# FunSPU: a versatile and adaptive multiple functional annotation-based association test of whole-genome sequencing data

**DOI:** 10.1101/350355

**Authors:** Yiding Ma, Peng Wei

## Abstract

Despite ongoing large-scale population-based whole-genome sequencing (WGS) projects such as the NIH NHLBI TOPMed program and the NHGRI Genome Sequencing Program, WGS-based association analysis of complex traits remains a tremendous challenge due to the large number of rare variants, many of which are non-trait-associated neutral variants. External biological knowledge, such as functional annotations based on ENCODE, may be helpful in distinguishing causal rare variants from neutral ones; however, each functional annotation can only provide certain aspects of the biological functions. Our knowledge for selecting informative annotations *a priori* is limited, and incorporating non-informative annotations will introduce noise and lose power. We propose FunSPU, a versatile and adaptive test that incorporates multiple biological annotations and is adaptive at both the annotation and variant levels and thus maintains high power even in the presence of noninformative annotations. In addition to extensive simulations, we illustrate our proposed test using the TWINSUK cohort (n=1,752) of UK10K WGS data based on six functional annotations: CADD, RegulomeDB, FunSeq, Funseq2, GERP++, and GenoSkyline. We identified genome-wide significant genetic loci on chromosome 19 near gene *TOMM40* and *APOC4-APOC2* associated with low-density lipoprotein (LDL), which are replicated in the UK10K ALSPAC cohort (n=1,497). These replicated LDL-associated loci were missed by existing rare variant association tests that either ignore external biological information or rely on a single source of biological knowledge. We have implemented the proposed test in an R package “FunSPU”.

## 1. Introduction

In recent years, large-scale whole-exome sequencing and whole-genome sequencing (WGS) data have been generated, such as those in the Exome Sequencing Project ^1^, the UK10K project ^2^ and the ongoing NIH NHLBI Trans-Omics for Precision Medicine (TOPMed) WGS program ^3^, providing unprecedented opportunities to investigate low-frequency variants (minor allele frequency [MAF] between 1% and 5%) and rare variants (RVs; MAF < 1%) in association with complex diseases and traits. However, WGS-based association analysis of complex traits remains a tremendous challenge due to the large number of RVs, many of which are non-trait-associated neutral variants. External biological knowledge, such as functional annotations, might be informative to distinguish causal RVs from neutral ones. Some recent large-scale functional genomic studies, such as ENCODE ^4^ NIH Epigenomics Roadmap ^5^ and GTEx ^6^ projects, provide rich resources to use in characterizing the functional consequences of single nucleotide variants (SNVs), especially those in non-coding regions. Many approaches have been developed for functional annotations by integrating these data, e.g., CADD ^7^, GenoSkyline ^8^ and Eigen ^9^; see Liu et al for a recent comparative review ^10^. In WGS analysis, investigators may filter a subset of SNVs by annotations ^2; 11^, or use a single source of functional scores as weights in association tests to boost the statistical power ^12–14^; however, each functional annotation can only provide a certain aspect of the biological functions, e.g., sequence conservation across species or biochemical activity of non-coding regions in a tissue. Our *a priori* knowledge to select the informative annotation(s) regarding a phenotype and genomic regions of interest is limited, and incorporating noninformative annotations will introduce noise and lose power.

To address this analytical challenge, we propose a family of versatile and powerful tests called “FunSPU” that allow for incorporating multiple <u>fun</u>ctional annotations simultaneously in the adaptive sum of powered score (a<u>SPU</u>) test framework ^15^. The fundamental idea of aSPU is to construct a general class of association tests, each of which is the most powerful under varying, yet unknown, local genetic architecture, then data-adaptively select the most significant test. Since each functional annotation system contains limited biological knowledge, multiple sources of functional annotations may provide complementary information. Therefore, a test that integrates multiple functional annotations simultaneously is potentially powerful. The proposed test is adaptive at both the annotation and variant levels and thus maintains high power even in the presence of noninformative annotations and a large number of neutral RVs. We also propose minimum *p*-value (minP) and Fisher’s meta-analysis-like approaches to combine the *p*-values with respect to multiple annotations. Moreover, to further increase the statistical power, we propose to incorporate a trait-specific global weight for each annotation based on partitioning the heritability.

Using extensive simulations and application to the UK10K WGS data ^2^, we compared our proposed FunSPU tests with the corresponding annotation-ignorant aSPU test as well as some existing RV association tests, such as the T5 burden test and SKAT ^16^. We also compared our method with a recently published multiple functional annotation-based association test called functional score test (FST) ^17^ Using the UK10K TWINSUK WGS cohort as the discovery sample (n=1,752), we considered six functional annotations, CADD ^7^, RegulomeDB ^18^, FunSeq ^19^, Funseq2 ^20^, GERP++ ^21^ and GenoSkyline ^8^, and four quantitative traits, low-density lipoprotein (LDL), high-density lipoprotein (HDL), body mass index (BMI) and systolic blood pressure (SBP). We identified genome-wide significant genetic loci on chromosome 19 near gene *TOMM40* [MIM: 608061] and *APOC4-APOC2* [MIM: 600745] that are associated with LDL, which are replicated in the UK10K ALSPAC WGS cohort (n=1,497). These replicated LDL-associated loci were missed by existing RV association tests that either ignore external biological information or rely on a single source of biological knowledge. We have implemented the proposed test in an R package “FunSPU”.

## 2. Methods

### 2.1. Notations

For the purpose of exposition, we introduce our proposed tests in the linear model framework with a quantitative trait and no covariates, though the methods can be similarly extended to binary or survival traits, and adjusted for covariates. Suppose that for subject *i* = 1,…, n, ***Y*** = (*Y*_*1*_,…, *Y*_*n*_) is the vector of a quantitative trait, and X_i_ = (*X*_*i1*_,…, *X*_*ik*_)’ is the vector of the genotype scores of *k* RVs, for example, from a gene or some genomic region. Here, we use additive coding for each RV; that is, *X*_*ij*_ is the count of the minor allele at RV *j* for subject *i*. For simplicity, we ignore other covariates in our model. We consider a linear regression model:

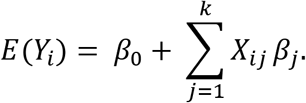

We test the null hypothesis *H*_0_; *β* = (*β*_*1*_,…, *β*_*k*_)’ = 0, that is, there is no association between any of the RVs and the trait under *H*_0_. Our proposed tests are based on the score vector *U* = (*U*_*1*_…, *U*_*k*_)’ for *β* and its covariance matrix *V*,

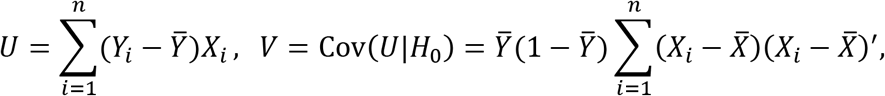

where *Y̅* and *X̅* are the sample means of the *Y*_i_’s and *X*_i_’s, respectively.

### 2.2. Review of the data-adaptive aSPU test

Pan et al. ^15^ proposed a new adaptive test that retains high power across a wide range of varying, yet unknown, genetic architecture for the analysis of RVs. This test is based on a class of the SPU test;

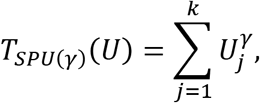

where γ≥1 is a positive integer. Suppose that we have a set of candidate values of γ in Γ, e.g., Γ = {1, 2, 3,…, 8,∞}, as used in our later experiments. It is known that SPU(1) is equivalent to the burden test, while SPU(2) is a variance-component score test equivalent to SKAT with a linear kernel. Importantly, as γ increases (as an even integer), the *SPU*(*γ*) test puts more weights on the larger component of *U* while gradually ignoring the remaining component. In particular, we have 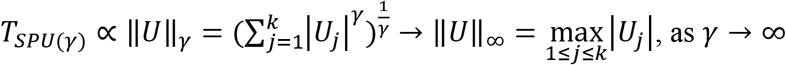. The SPU(∞) is closely related to the minP test (but ignores possibly varying variances of *U*_*j*_’s); the two tests often perform similarly ^22^ Since the power of an *SPU*(*γ*) test depends on the choice of *γ* while the optimal choice of *γ* depends on the unknown true association pattern of the RVs to be tested, it would be desirable to data-adaptively choose the value of *γ*. To this end, the aSPU test takes the minimum *p*-value of the SPU(γ) tests as its test statistic: 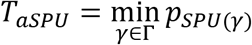. In this case, *T*_*aSPU*_ is no longer a genuine *p*-value; we use resampling approaches such as residual permutation or parametric bootstrap to obtain its *p*-value.

### 2.3. New test: FunSPU - a data-adaptive test incorporating multiple annotations

Our proposed test is in the data-adaptive aSPU test framework. Importantly, the proposed test is adaptive at both the annotation and SNV levels. Suppose that we have the score vector *U* = (*U*_*1*_…, *U*_*k*_)’ for *k* RVs from a gene region or sliding window based on a linear regression model. Let 0 ≤ *w*_*lj*_ ≤ 1 denote the functional score from the *l*th of *m* properly scaled annotations for the *j*th of *k* RVs. The proposed functional annotation-based SPU test is

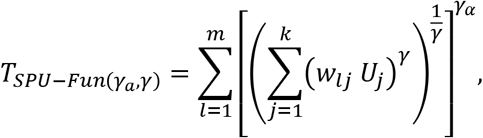

where two positive integers γ ≥ 1 and γ_*a*_ ≥ 1 respectively control the individual variants’ and annotations’ relative contributions to the overall test statistic; e.g., γ_a_ = 1 treats all annotations equally, while γ_a_ = ∞ only considers the most significant annotation. The inner sum of weighted *U*_*j*_ with power γ is the weighted SPU, and they are normalized to the power of 1/γ before being subjected to the outer sum with power γ_a_. Since the number of the RVs in this test statistic is identical across all *m* annotations, it is not necessary to further normalize the weighted SPU test by the number of RVs.

The intuition to use γ_a_ as the powers of the weighted SPU is similar to that for γ. In general, a smaller γ_a_, e.g., γ_a_ = 1, is more effective when there are more informative annotations, each of which is roughly equally discriminative regarding the deleteriousness of the RVs for the trait of interest. In contrast, a larger γ_a_ is preferred if there is only one or fewer informative annotations that can well distinguish causal variants from neutral ones for the trait. As γ_a_ → ∞, only the most significant weighted SPU is considered.

We aim to perform powerful tests when there are unknown association patterns of RVs and unknown informativeness of functional annotations. In practice, since we have no *a priori* knowledge about choosing γ and γ_*a*_, we need to conduct a grid search over a set of possible values of both γ and γ_*a*_. However, searching too many values will introduce extra variability and lead to reduced power. This effect was later confirmed when we used γ_*a*_ ∈ {1,2,3,…, 8, ∞} and γ ∈ {1,2,3,…, 8, ∞} in some preliminary simulations. Based on the results of aSPU tests ^15^ and the feature of annotations, we decided to use γ_*a*_ ∈ Γ_*a*_ = {1,2,4,8, ∞} and γ ∈ Γ = {1,2,3,…,6} for the rest of the study. We retained γ_*a*_ = ∞ as an approximation to the minP test and ignored some higher values of γ since the results tend to be similar to γ = 6.

Given a set of γ and γ_*a*_, e.g., γ ∈ Γ = {1,2,3,…,6} and γ_*a*_ ∈ Γ_*a*_ = {1,2,4,8, ∞}, the proposed data-adaptive FunSPU test statistic is defined as

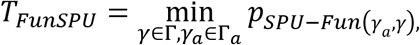

where 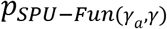 is calculated by the resampling methods detailed below. Although the score vector U has an asymptotic normal distribution N(0, V), it is not easy to derive the asymptotic distribution of *T*_*FunSPU*_. Therefore, we propose using a single layer of permutations (without covariates) or residual permutations (with covariates) to obtain *p*-values as done in aSPU ^15; 23^. Specifically, we first permute the original set of trait *Y* to obtain a new set of *Y*^*(b)*^ for *b* = 1,…, *B*. Then, we calculate the null score vector *U*^*(b)*^ and the corresponding test statistic 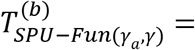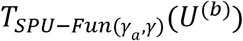 as well as their *p*-values values 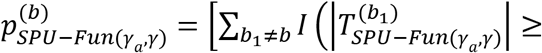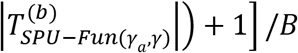 Therefore, we have 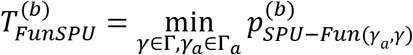, and the final *p*-value of the FunSPU test 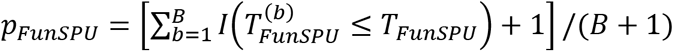.

In the FunSPU test above, we ignored the possibly different variances of the score function component *U*_*j*_, for example, due to varying MAF of the RVs. On the other hand, previous research has shown that it may be beneficial to account for the heterogeneity of variances in the SPU framework ^22^ Therefore, we further propose an inverse-variance weighted version of FunSPU:

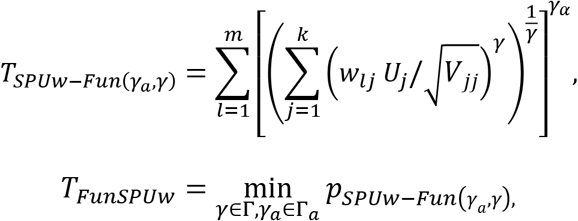

where *V*_*jj*_ is the *j*th diagonal element of *V* = Cov(*U*|*H*_0_) as given before.

### 2.4. Alternative approaches to incorporating multiple functional annotations: aSPU_minP and aSPU_Fisher

We considered alternative approaches to incorporate multiple functional annotations into the aSPU test. In contrast to the two-level FunSPU approach, we can obtain modified aSPU tests via the score vector *U* weighted by each functional annotation, i.e., 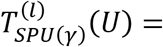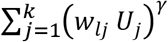 and 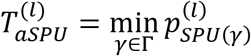, for *l* = 1,…, m. We can obtain the genuine *p*-value 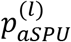 by resampling methods. To combine multiple functional annotations, we can further employ some general approaches to combine multiple *p*-values, 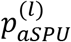. For example, we can simply use 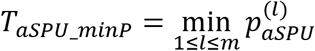 as the test statistic of *m* modified aSPU tests. This aSPU_minP test is similar, but not exactly equivalent to the case of FunSPU with γ_*a*_ = ∞; the latter chooses the maximum 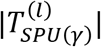 and then uses resampling methods to obtain a genuine *p*-value directly, while the aSPU_minP test calculates the empirical *p*-value 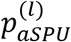 first, and then uses the minimum *p*-value *T*_*aSPU_minP*_ as the new test statistic and resampling to calculate the final *p*-value.

Another common method for combining *p*-values is Fisher’s meta-analysis approach, i.e., 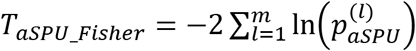. If the *m p*-values were independent, *T*_*aSPU_Fisher*_ would follow a chi-squared distribution with 2*m* degrees of freedom. However, our 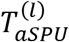 tests are correlated via the score vector *U*. Hence, we also use resampling approaches to calculate the final *p*-value. We can similarly apply the inverse-variance weighted method to aSPU_minP and aSPU_Fisher tests, respectively denoted as aSPUw_minP and aSPUs_Fisher.

### 2.5. wtFunSPU: extension of FunSPU to allow for global weighting of multiple annotations

In our proposed FunSPU test, we treated all *m* functional annotations equally *a priori* and completely relied on the data to adaptively combine multiple annotations in each test unit, for example, a sliding window. This may be less efficient in the presence of overall inferior or superior annotations for a trait of interest, in which case it would be desirable to globally down-weight inferior annotations (and up-weight superior annotations). To this end, we propose to modify the FunSPU test by introducing an annotation-level weight *ρ* = (*ρ*_1_,…,*ρ*_*m*_)′ and denote the modified test as wtFunSPU;

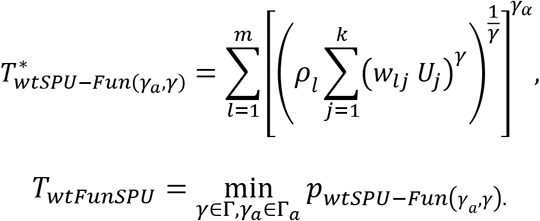

Since we assume no *a priori* knowledge regarding the informativeness of a functional annotation for a given trait, we propose to estimate *ρ*_*l*_ based on some global correlation measure between the annotation weights, genotypes and phenotype. A promising approach is based on partitioning the heritability *h*^2^ by functional annotations ^24^: a functional annotation is more informative for the trait of interest if SNVs with higher functional scores contribute to more heritability on average. Specifically, given an annotation, we first partition the genome-wide RVs based on Q discrete functional categories or percentiles of continuous functional scores; we then estimate the heritability 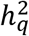 for all SNVs in functional category *q* = *1,…, Q*, using the GCTA tool ^25^. We next compute the average per-SNV heritability 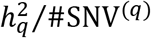 for each annotation category *q* and regress 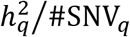 on q to estimate the slope: 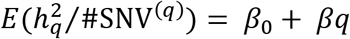, where *β* is used as the global weight *ρ* for the corresponding functional annotation in the wtFunSPU test. Prior to this calculation, we transform the functional annotation to positive integers *q* = *1,…, Q* such that larger *q* corresponds to a more likely functional category. If a functional category has a very small number of SNVs or 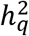 close to zero, this category is combined with a nearby category; see Supplemental Figures S1 to S4 and Table S1 for details.

## 3. Results

### 3.1. Simulation setups

We conducted extensive simulations to evaluate and compare the performance of our proposed functional annotation-based tests with existing association tests for RVs. To make the simulation study representative of real RV data, we randomly selected 200 RVs from chr16:56.8M~57.1M of the UK10K TWINSUK genotype data of 1,718 unrelated individuals. MAFs of the selected RVs were no larger than 1%.

To evaluate power, we generated the simulated phenotypes as follows. First, we simulated 3 sets of informative annotations (*w*_1*j*_, *w*_2*j*_,*w*_3*j*_) and 3 sets of random annotations (*w*_4*j*_, *w*_5*j*_,*w*_6*j*_) independently(*j* = 1, 2,…, 200 ordered by genomic positions). We designated the first 100 RVs as causal variants (*j* = 1, 2,…, 100) and the remaining 100 RVs as neutral variants (*j* = 101, 102,…, 200). The informative annotations were generated from a uniform distribution *U*(0.4, 1) corresponding to causal variants and from *U*(0, 0.6) corresponding to neutral variants. All of the random annotations were generated from *U*(0, 1). Second, we randomly selected *k* = *k*_1_+ *k*_2_ RVs: *k*_1_ causal RVs from *j* = 1, 2,…, 100 and *k*_2_ neutral RVs from *j* = 101, 102,…, 200. Third, we used only informative annotations to calculate the effect size = (*w*_1*j*_, *w*_2*j*_,*w*_3*j*_) for each causal RV. Fourth, the simulated phenotype was obtained from 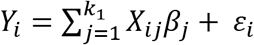, where *ɛ*_*i*_ followed *N*(0,3) and *i*=1,2,…,1718. Furthermore, to evaluate the globally weighted wtFunSPU test, we calculated the correlations between the sum of the genotypes weighted by each annotation and simulated phenotypes for each of the 1,000 simulation replications, and used the mean of the 1,000 correlations as the global weight of each annotation, i.e., 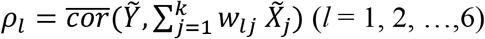, where *Ỹ* and *X̃*_*j*_ are the vectors of *Y*_*i*_ and *X*_*ij*_ in each replication correspondingly

We considered two simulation scenarios. In scenario A, we used all three informative annotations, three random annotations and one dummy annotation (1’s for all RVs) in functional annotation-based tests (FunSPU, aSPU_minP, and others). To test the effect of more “noisy” annotations, we implemented scenario B, which used only one informative annotation, all three random annotations and one dummy annotation in the tests. In both scenarios A and B, we used identical procedure as above to generate simulated phenotypes *Y*_*i*_, and fixed *k*_1_ = 8 and *k*_2_ = {8, 16, 32, 64, 128}, respectively. We set *C*_*β*_ =0.5 for tests that incorporated global weights and *c*_*β*_ =1 for other tests.

To evaluate the type I error rate, we simulated *Y*_*i*_~*N*(0,3) (*i*=1,2,…,1718), independent of *k* neutral RVs and 6 random annotations all from *U*(0,0.6) in each replication. We set increasing numbers of neutral RVs with *k*={8, 16, 32, 64, 128}.

Throughout the simulations, we fixed the significance level at α = 0.05. The empirical power was calculated based on 1,000 replications for each scenario, and the empirical type I error rate was calculated based on 5,000 replications. For permutation-based tests, 1,000 resamplings were conducted for each replication.

### 3.2. Simulation results

As shown in Table 1, all the tests under comparison could control the type I error rate satisfactorily around 0.05. Regarding power, we first considered scenario A (Figure 1), which was an advantageous scenario for our proposed tests since all three informative annotations together with three random annotations and one dummy annotation were used in the tests. The dummy annotation (constant 1) was supposed to retain the unweighted SPU in the adaptive tests, as in aSPU. Although the simulated annotations for causal and neutral RVs had modest differences, i.e., from *U*(0.4, 1) and *U*(0,0.6), respectively, the tests incorporating functional annotations, such as FunSPU, wtFunSPU and aSPU_minP, always had higher power than tests that ignored functional annotations, such as aSPU, SKAT and T1. The FunSPU test appeared to be less powerful than aSPU_minP, suggesting a lack of efficiency in the former’s complete data-adaptive strategy to combine multiple annotations. On the other hand, wtFunSPU and wtFunSPUw outperformed aSPU_minP, supporting the effectiveness of the global weighting scheme. We also observed that the inverse-variance weighted tests always outperformed the original tests, e.g., wtFunSPUw versus wtFunSPU, and this advantage became more obvious with a higher proportion of neutral RVs. Lastly, the power of the aSPU_Fisher test was similar to that of the aSPU_minP test (results not shown).

**Table 1.**
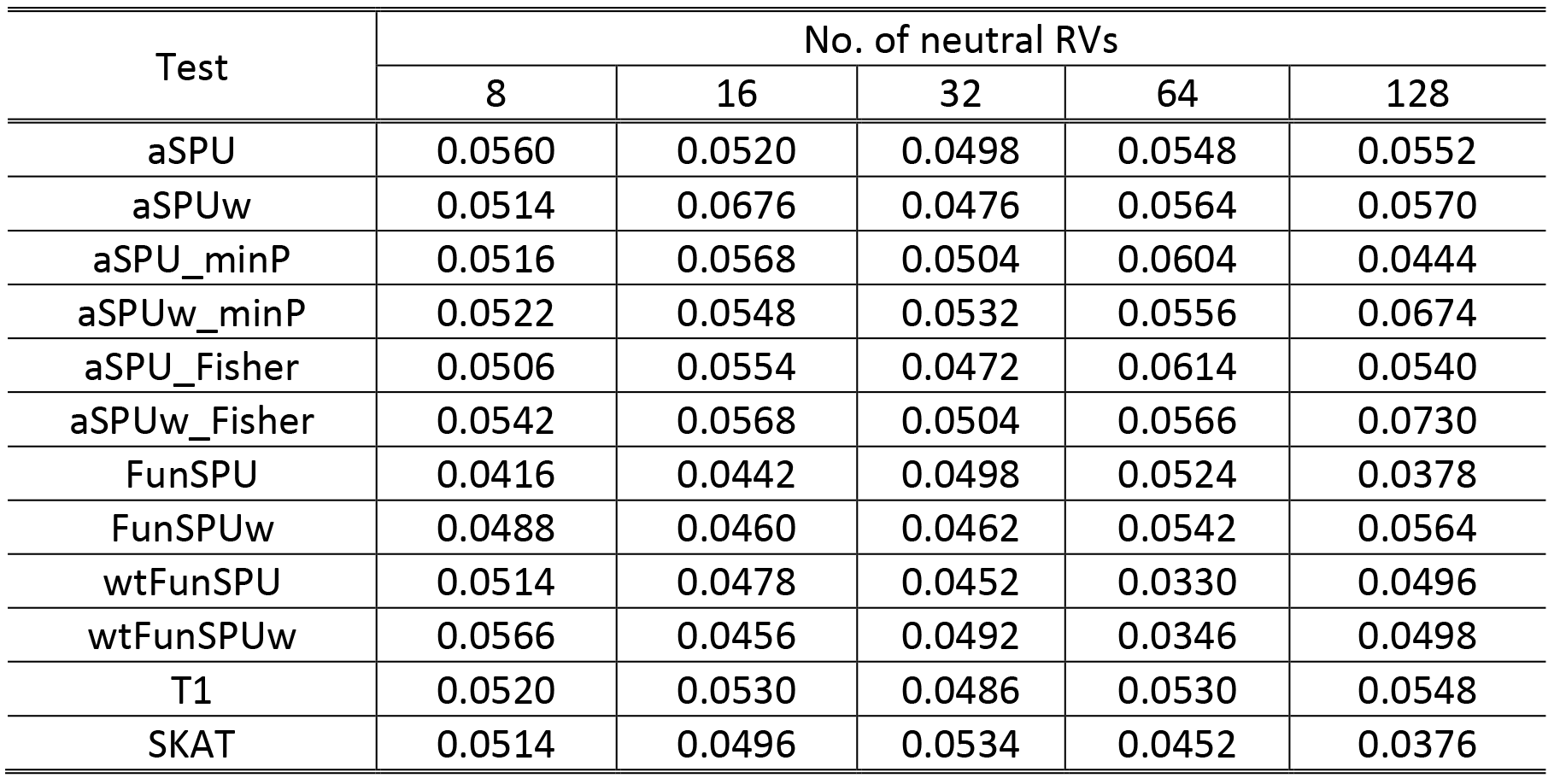
Empirical type I error rates of various tests for increasing number of neutral RVs with 5,000 simulation replications. Annotation-based tests were based on six random annotations.

**Figure 1.**
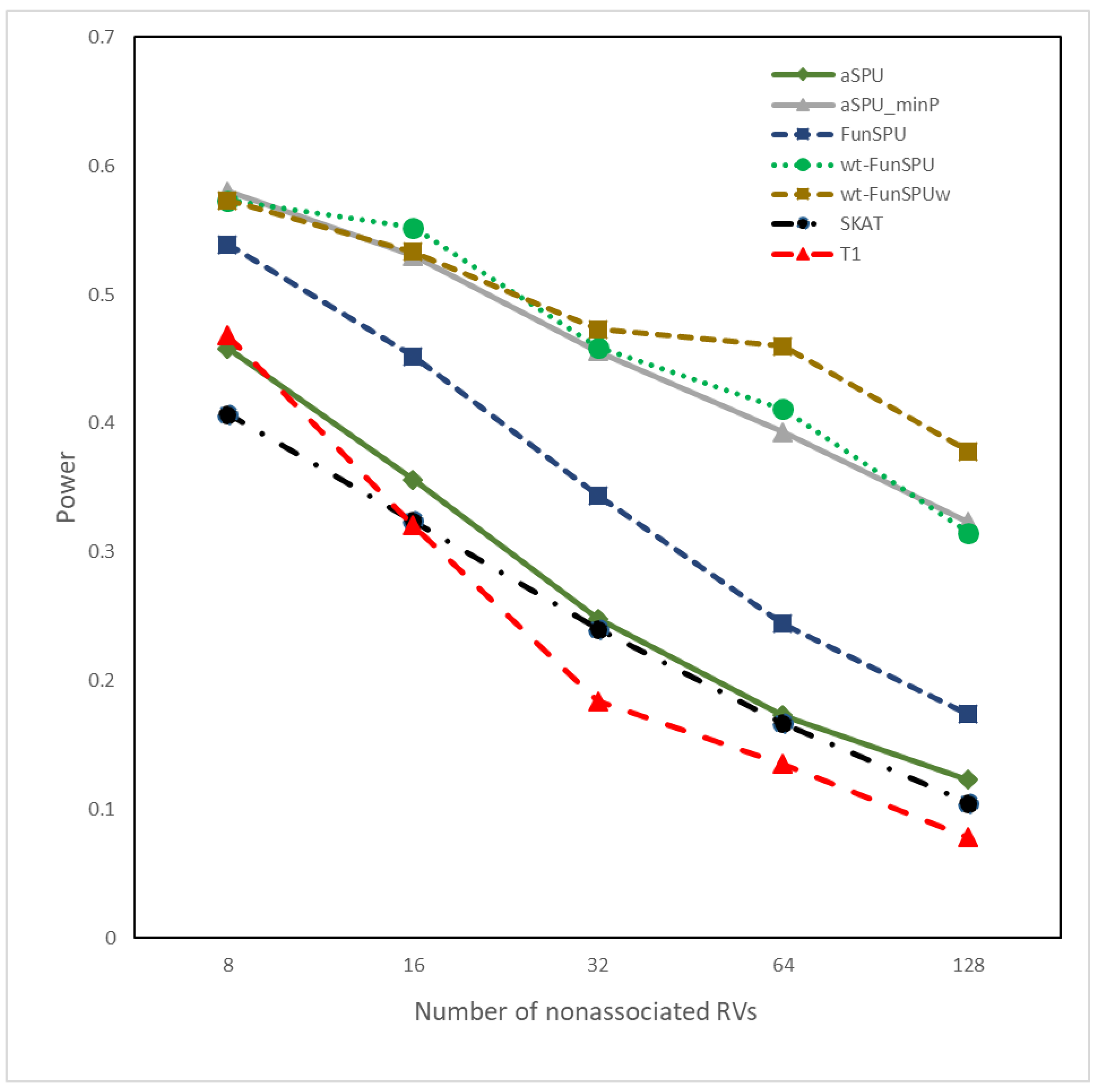
Empirical power of various tests for eight causal RVs and increasing number of nonassociated RVs. The incorporated annotations for association tests include all three informative annotations and three noninformative annotations (Scenario A). All the results were based on 1000 simulation replications.

Next, we considered a weaker scenario for our proposed tests. In scenario B (Figure 2), we used only one informative annotation, but all three random annotations and one dummy annotation in the tests. In this case, we had a higher proportion of “noisy” annotations in our tests. We observed that the FunSPU test was marginally more powerful than aSPU, SKAT and T1, but was less powerful than the aSPU_minP test by a large margin. In fact, scenario B was an advantageous scenario for the latter test, which only considered the most informative annotation. Finally, the globally weighted wtFunSPU and wtFunSPUw, especially the latter, were more powerful than the aSPU_minP test, again suggesting the benefit of globally down-weighting noninformative annotations.

**Figure 2.**
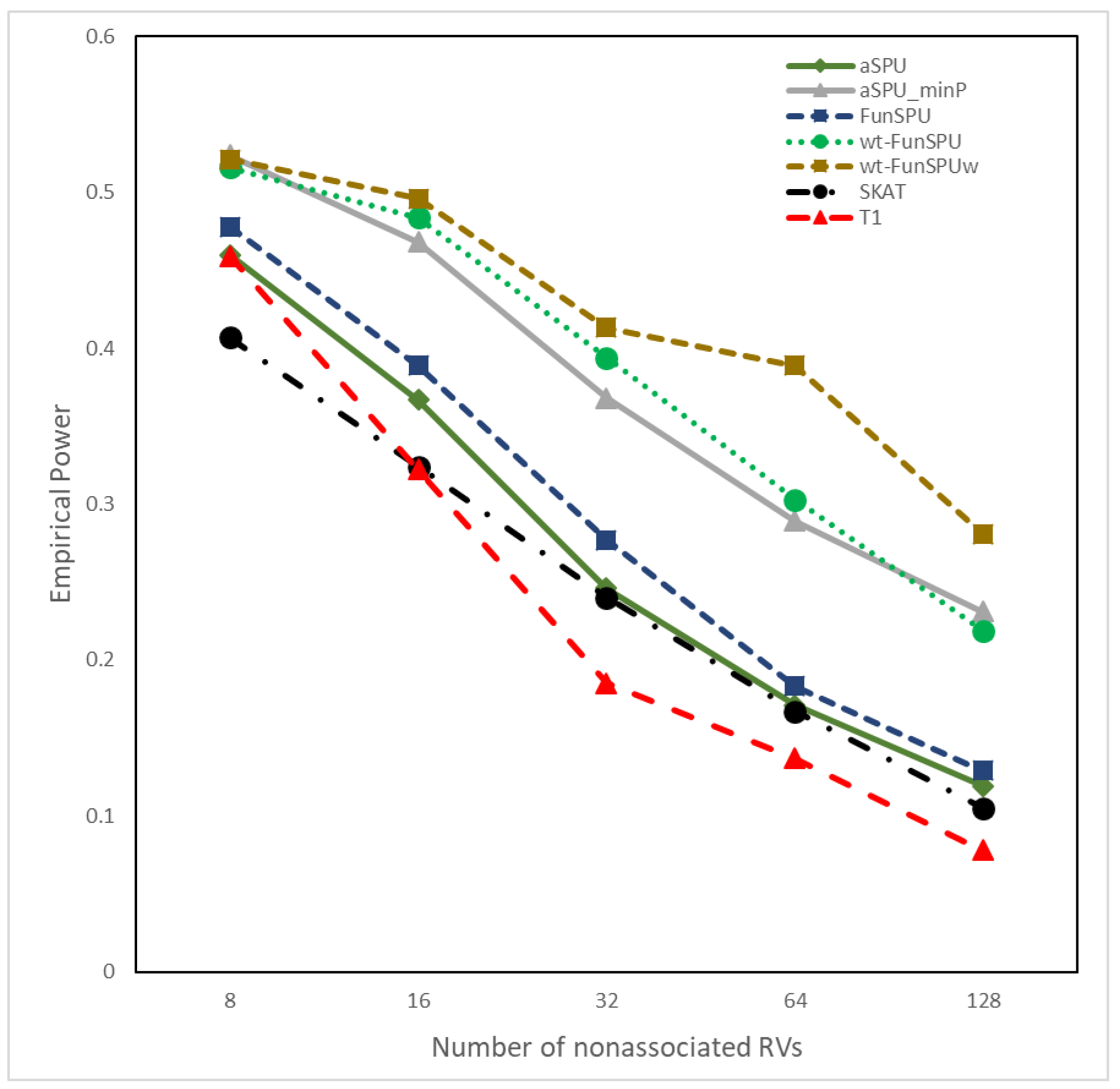
Empirical power of various tests for eight causal RVs and increasing number of nonassociated RVs. The incorporated annotations for association tests include one out of three informative annotations and three noninformative annotations (Scenario B). All the results were based on 1000 simulation replications.

### 3.3. Application to the UK10K WGS data

To further evaluate the performance of our proposed tests on real data, we applied FunSPU and other state-of-the-art tests, including SKAT, T5 burden test and FST (combined test)^17^, to association analysis of the UK10K WGS data with four complex quantitative traits: LDL, HDL, BMI and SBP. We used the TWINSUK samples as the discovery cohort and the ALSPAC samples as the replication cohort with n=1706/1497 (TWINSUK/ ALSPAC), 1718/1497, 1752/1792 and 1740/1796, respectively, for LDL, HDL, BMI and SBP, after merging WGS genotype and phenotype data. After removing SNVs that did not pass quality control as done in the original UK10K analysis^2^, as well as singletons and INDELs, we had a total of 10,979,027 RVs and low-frequency variants with MAF< 5% in the discovery cohort.

We considered six types of functional annotations for RVs. CADD^7^, FunSeq^19^, FunSeq2^20^, RegulomeDB^18^ and GERP++^21^ were extracted from the precomputed WGSA ^26^ library, and GenoSkyline (blood) annotation was generated from the region-based GenoSkyline library ^8^. We re-scaled all annotations to numerical weights within the interval (0, 1), with larger weights corresponding to a greater likelihood of being functional (Supplemental Figure S5). Among the above annotations, CADD, Funseq2, GenoSkyline and GERP++ provided rank scores, requiring no transformation. The RegulomeDB categories *s* = (1,2,…, 6) were transformed into (0, 1) by f(s)=(7-s)/6, where the Funseq categories *s* = (0,1,2,…, 6) were transformed by f(s)=(1+s)/7. We substituted the missing values or zero values with 0.01 (FunSeq, FunSeq2, RegulomeDB) or 0.0001 (GERP++). There was no missing value in CADD and GenoSkyline for the RVs considered here. Supplemental Figure S6 shows the pairwise correlation coefficients among the 6 annotations: while some annotations were moderately correlated (r > 0.3), for example, GERP++ with CADD, and Funseq2 with RegulomeDB/Genoskyline, others were much less correlated. This suggests that multiple annotations may provide complementary information regarding the functional consequence of genetic variants, and it may be beneficial to incorporate them simultaneously in association analysis as proposed in the FunSPU framework here. Following the procedure proposed in Section 2.5, we calculated the phenotype-specific weight for each of the six annotations and used them as global weights in the wtFunSPU test. As shown in Supplemental Figures S1 to S4 and Table S1, RegulomeDB, Funseq and GenoSkyline tended to have consistently higher weights than GERP++, Funseq2 and CADD, while the numerical values and the relative magnitudes of the weights could vary across phenotypes.

We employed a sliding window approach to group RVs with a window length of 10k base pairs (bp) and a step size of 8.75k bp, resulting in 319,306 windows in total. Using the conservative Bonferroni procedure, we set the family-wise error rate at 0.05 with a significance level = 0.05/319306 = 1.56e-07, which equals 6.81 on the log_10_ scale. To achieve this genome-wide significance level, we used a step-up permutation strategy ^23; 27^ We first performed *B*=10,000 permutations for all sliding windows and gradually increased *B*; if those sliding windows with estimated *p*-values <10/B, we increased *B* to 10 times the current value and reestimated the *p*-values for these sliding windows. The number of permutations in the final stage was *B*=10^8^.

As shown in Supplemental Figures S7 to S10, the quantile-quantile plots for the proposed FunSPU tests were well behaved, with no discernible indication of global *p*-value inflation, suggesting that the FunSPU tests could control the type I error rate well in genome-wide scans. Table 2 shows all sliding windows with at least one genome-wide significant *p*-value in the TWINSUK discovery cohort by any of the association tests under consideration. In total, the global weighted wtFunSPU and wtFunSPUw tests identified 14 genome-wide significant sliding windows, whereas the FunSPU and FunSPUw tests identified 8 significant sliding windows, and the FST test identified 6 significant sliding windows, respectively. As a comparison, SKAT and T5 each only identified one significant sliding window, suggesting that incorporating functional annotations may improve the power of association analysis of WGS data, and our proposed adaptive tests are competitive compared with existing methods such as the FST.

**Table 2.** Genome-wide significant sliding windows identified by various tests in the UK10K TWINSUK cohort and replication in the ALSPAC cohort of UK10K. Significant *p*-values are in boldface; only significant *p*-values in the ALSPAC cohort were reported (TWINSUK *p*-value/ALSPAC *p*-value in shade). cMAF: cumulative minor allele frequency. Base pair (bp) position based on reference genome hg19.

| Trait | Chr | Start position–<br>Stop position (bp) | Gene(s) | cMAF | # SNVs | *p*-Values |  |  |  |  |  |  |  |  |  |
| --- | --- | --- | --- | --- | --- | --- | --- | --- | --- | --- | --- | --- | --- | --- | --- |
|  |  |  |  |  |  | Globally weighted |  | Not globally weighted |  | aSPU<br>_minP | aSPUw<br>_minP | aSPU | FST<br>(combined) | SKAT | T5 |
|  |  |  |  |  |  | wtFun<br>-SPU | wtFun<br>-SPUw | Fun-SPU | Fun-SPUw |  |  |  |  |  |  |
| HDL | 1 | 39,070,016–39,080,016 | Intergenic | 0.14/0.33 | 13/38 | 4.3e-2 | 2.3e-3 | 3.3e-2 | **<1e-8** | 2.0e-2 | 3.2e-6 | 3.6e-2 | 1.2e-3 | 5.2e-2 | 2.0e-4 |
| HDL | 2 | 173,903,159–173,913,159 | RAPGEF4 | 0.084/0.21 | 8/31 | **<1e-8** | **<1e-8** | 2.4e-6 | **<1e-8** | 4.9e-6 | 8.9e-6 | 2.4e-5 | 2.5e-6 | 1.1e-6 | 1.2e-4 |
| HDL | 3 | 63,804,000–63,814,000 | C3orf49 | 0.02/0.17 | 4/36 | 2.9e-3 | **<1e-8** | 4.0e-2 | 1.8e-5 | 3.1e-2 | 4.5e-5 | 0.12 | 3.2e-3 | 1.6e-2 | 5.2e-2 |
| HDL | 5 | 6,548,648–6,558,648 | Intergenic | 0.001/0.32 | 2/59 | 0.13 | **<1e-8** | 0.18 | **<1e-8** | 6.2e-2/ **<1e-4** | **<1e-8/ <1e-4** | 0.15 | 2.0e-5 | 2.1e-3 | 0.19 |
| HDL | 5 | 6,557,398–6,567,398 | Intergenic | 0.34/0.26 | 42/43 | 4.3e-2 | 1.5e-2 | 0.13 | **<1e-8** | 9.5e-3 | **<1e-8** | 0.24 | **1.2e-8** | 1.8e-2 | 0.81 |
| LDL | 3 | 102,287,400–102,297,400 | Intergenic | 0.25/0.19 | 24/39 | 4.4e-5 | 9.9e-3 | **<1e-8** | 3.6e-2 | 9.0e-6 | 1.3e-2 | 8.1e-6 | 1.3e-4 | 2.5e-5 | 4.5e-3 |
| LDL | 3 | 102,427,400–102,437,400 | Intergenic | 0.16/0.27 | 23/40 | **<1e-8** | 5e-8 | **<1e-8** | 1.3e-3 | 1.2e-6/ **<1e-4** | 5.1e-5 | 2.5e-4 | 3.3e-7 | 1.3e-4 | 0.68 |
| LDL | 5 | 43,259,958–43,269,958 | NIM1K | 0.43/0.29 | 37/46 | **9e-8** | **<1e-8** | 6.5e-7 | 3.0e-7 | 1.7e-5/ **<1e-4** | 8.2e-6 | 2.5e-5 | 1.5e-5 | 2.4e-3 | 4.4e-6 |
| LDL | 12 | 13,771,517–13,781,517 | GRIN2B | 0.18/0.23 | 43/161 | 3.5e-3 | 5.8e-4 | 0.80 | 3.5e-5 | 5.9e-6 | 1.7e-6 | 0.60 | **2.4e-11** | 0.23 | 0.96 |
| LDL | 12 | 13,780,267–13,790,267 | GRIN2B | 0.26/0.27 | 45/140 | 6.0e-2 | 8.0e-4 | 0.47 | 1.3e-4 | 1.0e-6 | 1.7e-6 | 0.33 | **2.0e-11** | 4.9e-2 | 0.53 |
| LDL | 19 | 45,387,096–45,397,096 | PVRL2/<br>TOMM40 | 0.21/ 0.22 | 33/ 37 | 5.4e-2/<br>**1.8e-3** | 0.10 | 3.5e-7/<br>**<1e-4** | 5.0e-3/<br>**1.6e-3** | 2.1e-7/ **<1e-4** | 4.5e-3 | **5.0e-8/ <1e-4** | 2.4e-4 | 2.4e-4/<br>**7.9e-6** | 0.25 |
| LDL | 19 | 45,395,846–45,405,846 | TOMM40 | 0.42/0.59 | 27/62 | 8.6e-3/<br>**<1e-4** | 8.6e-2 | **3.0e-8/ <1e-4** | 1.4e-4/<br>**1.9e-3** | **<1e-8/ <1e-4** | 5.8e-5/ **<1e-4** | 5.0e-7 | 1.2e-4/<br>**1.0e-10** | 4.7e-4/<br>**1.1e-6** | 0.28 |
| LDL | 19 | 45,439,596–45,449,596 | APOC4-APOC2 | 0.37/0.18 | 22/25 | **<1e-8/ <1e-4** | 1.1e-4 | 1.2e-6/<br>**<1e-4** | 6.9e-4 | 2.1e-5/ **<1e-4** | 1.4e-4 | 2.3e-4 | 4.7e-5/<br>**2.4e-7** | 1.1e-3/<br>**9.0e-4** | 0.15 |
| BMI | 3 | 35,619,294–35,629,294 | Intergenic | 0.24/0.24 | 16/40 | **<1e-8** | 4.2e-5 | 5.0e-5 | 1.1e-4 | 3.9e-5 | 3.6e-5 | 8.0e-5 | 2.9e-6 | 4.4e-5 | 1.5e-5 |
| BMI | 4 | 22,825,237–22,835,237 | Intergenic | 0.18/0.24 | 47/144 | 1.3e-3 | 5.2e-5 | 1.8e-3 | 6.0e-5 | 1.6e-4 | **3.4e-4/ <1e-4** | 2.8e-3 | **2.6e-8** | 1.4e-6 | 3.0e-5 |
| BMI | 10 | 13,937,041–13,947,041 | FRMD4A | 0.28/0.30 | 49/157 | 8.5e-2 | 2.4e-3 | 0.16 | 1.4e-4 | 5.9e-4 | 2.4e-4 | 0.18 | **1.9e-8** | 0.12 | 7.2e-2 |
| BMI | 12 | 26,179,017–26,189,017 | RASSF8 | 0.38/0.40 | 37/138 | 7.5e-2 | 1.2e-4 | 0.12 | 4.4e-4 | 2.1e-4 | **3.5e-4/ <1e-4** | 0.12 | **7.5e-8** | 6.6e-2 | 2.1e-4 |
| BMI | 15 | 42,935,528–42,945,528 | STARD9 | 0.28/0.23 | 26/56 | **<1e-8** | 2.7e-6 | **<1e-8** | 1.2e-5 | 2.1e-6 | 4.5e-5 | 1.2e-6 | 4.8e-7 | **8.4e-8** | **1.1e-7** |
| BMI | 16 | 60,159,304–60,169,304 | Intergenic | 0.27/0.12 | 20/28 | 4.3e-4 | **<1e-8** | 2.5e-2 | 2.1e-4 | **6.1e-4/ <1e-4** | **6.4e-6/ <1e-4** | 5.1e-3 | 1.4e-5 | 0.15 | 3.0e-4 |
| BMI | 21 | 40,851,354–40,861,354 | SH3BGR | 0.26/0.17 | 50/26 | 4.4e-4 | **<1e-8** | 2.5e-3 | 1.1e-4 | 1.3e-3 | **6.7e-5/ 2.6e-3** | 1.2e-3 | 3.9e-4 | 1.5e-2 | 3.4e-4 |
| SBP | 4 | 118,855,917–118,865,917 | Intergenic | 0.36/0.16 | 39/27 | 5.5e-2 | **<1e-8** | 0.68 | 8.1e-5 | 5.2e-2 | **4.2e-5/ <1e-4** | 0.75 | 2.5e-2 | 0.39 | 0.34 |
| SBP | 6 | 42,558,580–42,568,580 | UBR2 | 0.96/0.095 | 71/22 | **<1e-8** | 1.3e-4 | 3.4e-4 | 5.6e-4 | 1.1e-6 | 1.6e-4 | 3.9e-4 | 1.7e-6 | 2.7e-3 | 7.8e-5 |
| SBP | 6 | 121,241,282–121,251,282 | Intergenic | 0.50/0.23 | 65/46 | **<1e-8** | 1.8e-4 | 1.2e-4 | 1.4e-3 | 1.6e-4 | **2.1e-3/ <1e-4** | 5.5e-4 | 3.0e-4 | 1.1e-4 | 7.0e-4 |
| SBP | 11 | 38,236,659–38,246,659 | Intergenic | 0.60/0.48 | 86/61 | **<1e-8** | 1.4e-4 | 8.6e-5 | 6.4e-6 | 2.1e-5 | 1.9e-4 | 3.8e-3 | 4.4e-6 | 8.5e-3 | 0.36 |

Significance threshold: $p < 1.56 \times 10^{-7}$ for TWINSUK and $p < 2 \times 10^{-3}$ for ALSPAC

To confirm our findings in the TWINSUK cohort, we performed replication analysis of the genome-wide significant sliding windows in the ALSPAC cohort. As shown in Table 2, four sliding windows were replicated for the corresponding phenotypes and association tests with a replication *p*-value < 0.05/24 = 2e−3 based on the Bonferroni correction for 24 sliding windows: three by functional annotation-based wtFunSPU, FunSPU, aSPU_minP and aSPUw_minP tests and one by the aSPU test. In contrast, none of the 6 sliding windows identified by the FST test in the discovery cohort was replicated; neither did SKAT nor T5 replicate any sliding window.

Three of the four replicated sliding windows were close to each other on chromosome 19 around *TOMM40* and *APOC4-APOC2* genes. These loci have been previously identified and replicated to be associated with LDL by large-scale meta-analysis of GWAS common variants ^28–30^; however, this was the first time they were identified to harbor LDL-associated RVs with fewer than a couple of thousand samples, suggesting that the power of the FunSPU test was boosted by incorporating external biological knowledge.

We also looked into the effects of multiple annotations on the FunSPU tests. Although some high scores were observed around the *TOMM40* and *APOC4-APOC2* gene regions for Funseq2, Funseq, RegulomeDB and GenoSkyline (Figure 3), they did not appear to be obviously different from those scores outside these two loci. Figure 4 shows the association signals of selected tests in this genomic region, whereas Supplemental Figure S11 shows all individual annotation-based aSPU tests. As for *APOC4-APOC2*, three of the six annotations, namely, Funseq2, RegulomeDB and GenoSkyline (Supplemental Figure S11-E, H, I), positively contributed to the highly significant *p*-values of wtFunSPU and FunSPU (Supplemental Figure S11-A, B), although none of these individual annotation-based aSPU tests would reach the genome-wide significance threshold, demonstrating the benefit of integrating multiple functional annotations in the FunSPU framework. RegulomeDB and GenoSkyline also had higher global weights for LDL (Supplemental Table S1 (B)), which further boosted the *p*-value of the wtFunSPU test to the genome-wide significance level. As for *TOMM40*, Funseq2, CADD and GERP++ (Supplemental Figure S11-E,F,D) positively contributed to the genome-wide significance of FunSPU and aSPU_minP (Supplemental Figure S11-A and Table 2); whereas wtFunSPU missed this locus due to its low global weighting of these three annotations (Supplemental Table S1 (B)). This suggests that wtFunSPU, FunSPU and aSPU_minP may complement each other and may be used together in association analysis of WGS data.

**Figure 3.**
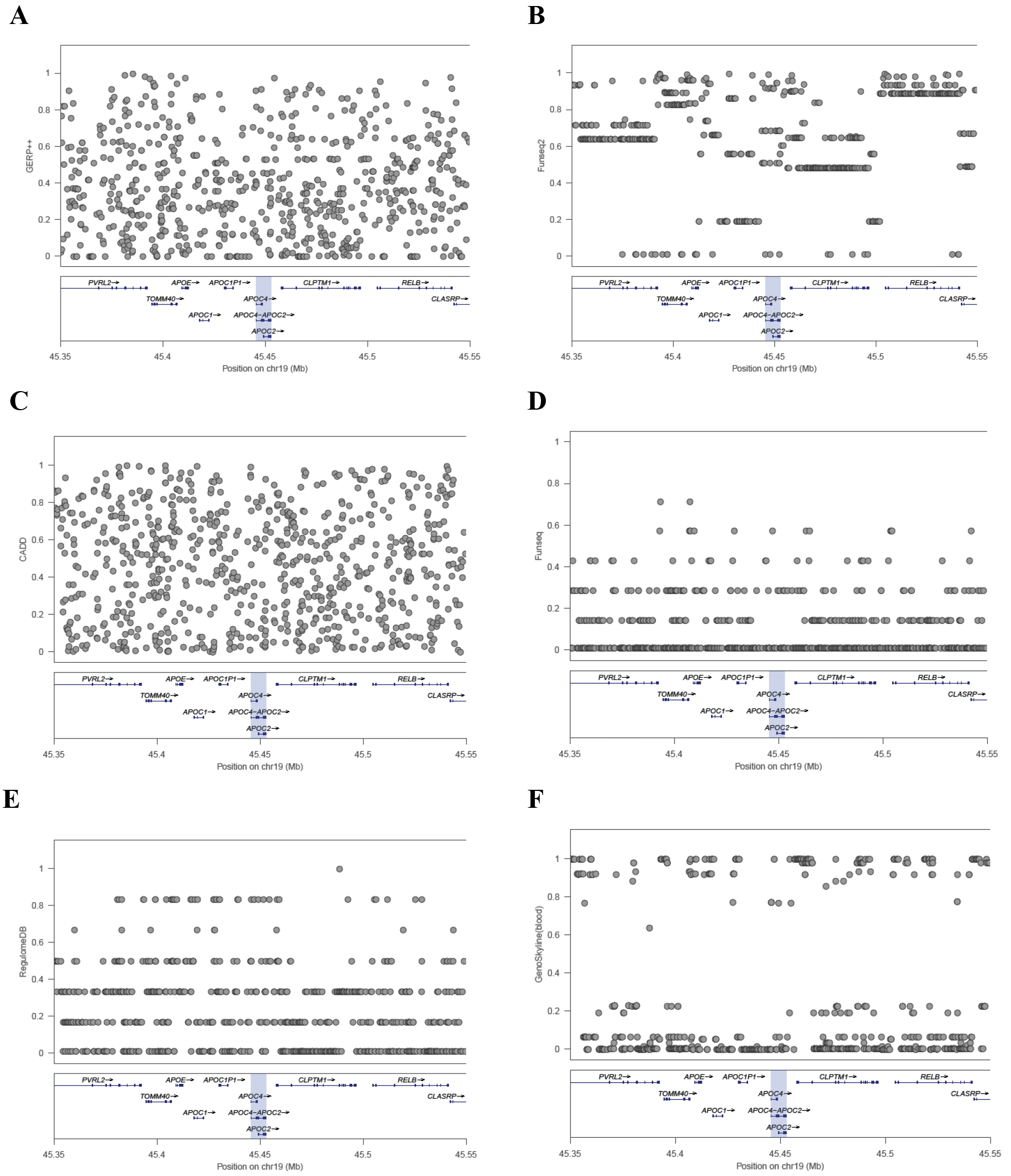
Rescaled scores of functional annotations: (A) GERP++, (B) Funseq2, (C) CADD, (D) Funseq, (E) RegulomeDB, and (F) GenoSkyline (blood) at the locus around gene *APOC4-APOC2*. The scores were rescaled to the interval [0, 1].

**Figure 4.**
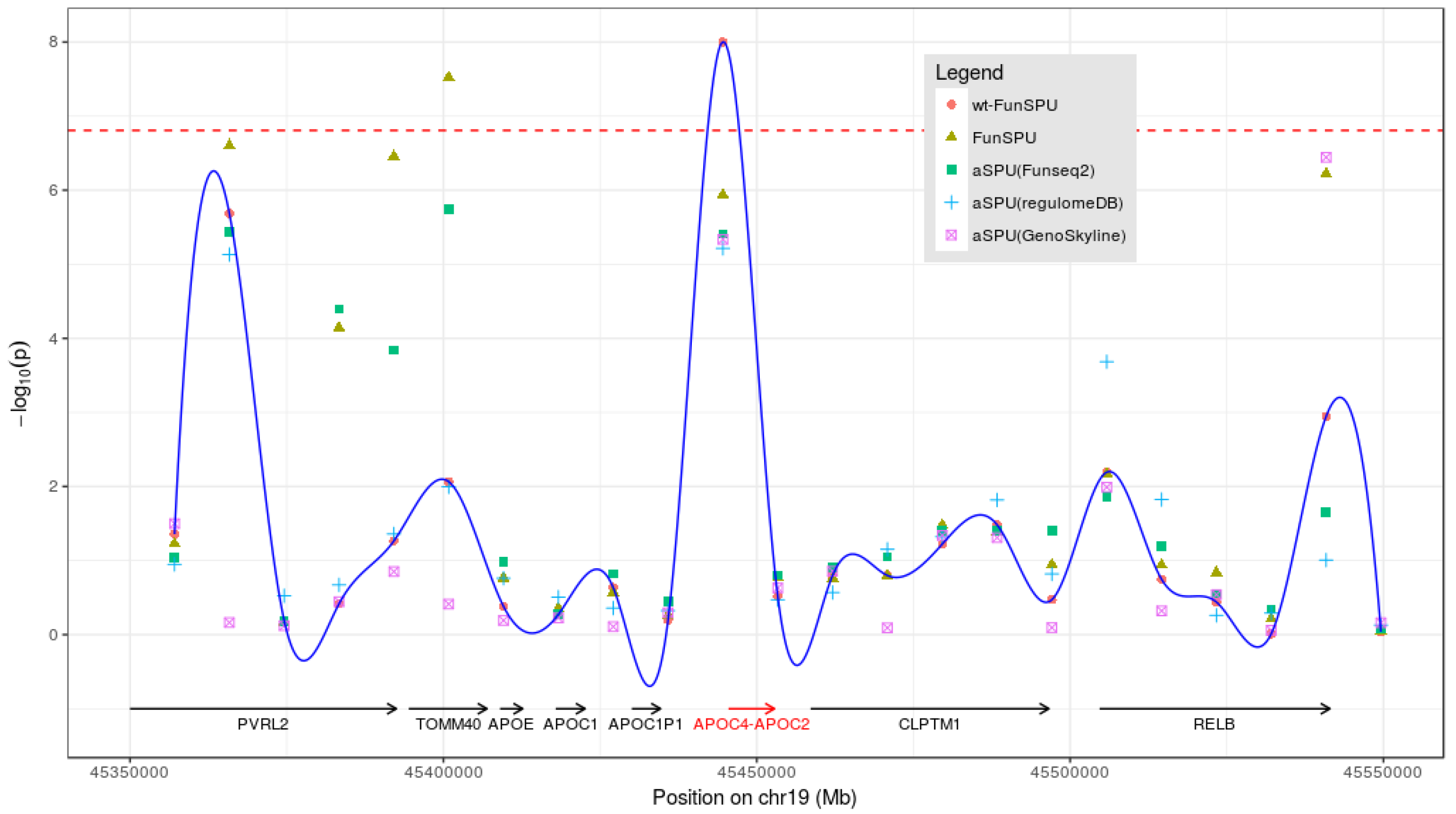
Association test results for LDL at the locus around gene *APOC4-APOC2*. The round points and the trace show the results from the globally weighted wtFunSPU test. Other points correspond to the results of FunSPU and single annotation-based aSPU (GERP++, Funseq2, RegulomeDB, GenoSkyline), respectively. Dashed line indicates the threshold of genome-wide significance level (*p* < 1.56e-7).

## Discussion

We have proposed a versatile and adaptive association test, FunSPU, to exploit multiple sources of biological knowledge in the analysis of WGS data. It is adaptive at both the annotation and variant levels, and thus maintains high statistical power, even in the presence of noninformative annotations and a larger number of neutral variants. We have further proposed a globally weighted wtFunSPU test to more effectively down-weight less informative functional annotations in a trait-specific manner. Using the UK10K WGS data, we demonstrated that our proposed FunSPU test and its extensions, including the wtFunSPU and aSPU_minP tests, are more powerful tools to identify genome-wide significant loci than existing RV association tests that either ignore external biological information or rely on a single source of biological knowledge. The FunSPU family of tests would thus serve as a powerful and complementary tool for ongoing and future large-scale WGS studies, such as the NHLBI TOPMed project^3^ of over 100,000 individuals and the UK Biobank^31^ WGS project of 50,000 individuals.

The six functional annotations we considered here are diverse in terms of resources and features. For example, GERP++^21^ is a sequence conservation score, whereas other annotations are ensemble scores based on integrating multiple sources of features, such as various functional genomic assays in the ENCODE project^4^ and eQTL evidence. As demonstrated in Supplemental Figure S6, a majority of the annotations were only moderately correlated with each other, supporting our proposal to incorporate multiple annotations’ approximately orthogonal yet complementary information regarding the functional consequence of RVs in the framework of the FunSPU association test. The FunSPU test can easily incorporate additional functional annotations, including some newly developed ones ^10^, such as fathmm-MKL^32^, Eigen/Eigen-PC^9^ and DeepSEA^33^.

To further de-noise noninformative annotations, we proposed a novel trait-specific measure based on partitioning the heritability and used it as a global weight for each annotation in the wtFunSPU test. Interestingly, our proposal is along the line of estimating group-specific weights in the context of weighted hypothesis testing ^34; 35^, though the latter is based on the mixture model, in contrast to the mixed model-based heritability partition here. Our proposed measure also has the potential to be used to compare the discriminative performance of whole-genome annotations for a complex trait of interest, for which known deleterious and neutral variants are rarely available (see Supplemental Table S1). This warrants further investigation.

We have some practical considerations for our proposed tests. First, some functional annotations are not well-defined across the whole genome, resulting in relatively high missing data rates, for example, 68% for Funseq; the missing scores may reduce the reliability of annotation-based association tests. On the other hand, considering multiple complementary functional annotations simultaneously may at least partially remedy the problem of missing information. Second, by employing parallel computing and a step-up residual permutation strategy for the FunSPU family of tests, we are able to perform computationally feasible genome-wide scans for WGS data. For example, in the UK10K TWINSUK WGS data application, it took 24 hours for 500 computing cores to complete the sliding window-based FunSPU scan, including 10e-8 residual permutations for the top sliding windows to reach the genome-wide significance threshold. We have implemented the proposed test in an R package “FunSPU”, available at https://github.com/sputnik1985/FunSPU; we expect that further implementation of the core functions in the C language should reduce the computational burden to a more affordable level.

## Supplemental Data

Supplemental Data include 11 figures and 1 table.

## Acknowledgments

This research was supported by National Institutes of Health grants R01HL116720, R01CA169122 and R21HL126032. We thank Drs. Wei Pan and Xiaoming Liu for helpful discussions and Ms. Lee Ann Chastain for editorial assistance. The authors acknowledge the Texas Advanced Computing Center at The University of Texas at Austin for providing HPC resources that have contributed to the research results reported within this paper. The authors declare no conflict of interest. This study makes use of data generated by the UK10K Consortium, derived from samples from the TwinsUK and ALSPAC cohorts. A full list of the investigators who contributed to the generation of the data is available from www.UK10K.org. Funding for UK10K was provided by the Wellcome Trust under award WT091310.

## Web Resources

FunSPU R package: https://github.com/sputnik1985/FunSPU

WGS Annotator (WGSA): https://sites.google.com/site/jpopgen/wgsa

OMIM: http://www.omim.org/

